# A murine model of adult gastrointestinal colonization by Group B *Streptococcus*

**DOI:** 10.1101/2025.09.25.678695

**Authors:** Joie Ling, Luke R. Joyce, Kelly S. Doran, Andrew J. Hryckowian

## Abstract

Group B *Streptococcus* (*Streptococcus agalactiae,* GBS) is a leading cause of invasive infections in neonates and adults. The adult gastrointestinal (GI) tract represents an understudied site of asymptomatic carriage with potential relevance for both transmission and disease. Here, we establish a murine model of GBS colonization in the adult GI tract, which provides a tractable system for probing host-microbe interactions within this niche. Using this model, we establish that GI carriage is generalizable to diverse GBS isolates and leverage transposon sequencing (Tn-Seq) to identify candidate GBS factors important for GI colonization. Informed by these TN-Seq data, we identify GBS capsule as a critical colonization factor of the adult murine GI tract. Taken together, this work highlights the GI tract as a reservoir for GBS and introduces a new experimental framework for investigating the bacterial and host determinants of GBS GI carriage.

## Introduction

Group B *Streptococcus* (*Streptococcus agalactiae*, GBS) is a disease-causing Gram-positive bacterial species. Found asymptomatically in the adult gastrointestinal (GI) tract and the female reproductive tract, GBS causes a wide variety of infections in neonates^1,2^, pregnant people^3^, and non-pregnant people^4–6^, especially immunocompromised adults and elderly adults. While preventative interventions decreased the incidence of GBS in neonates, the incidence of GBS disease in non-pregnant adults is increasing worldwide^7^. This highlights the limitations of current preventative measures and treatments for GBS and emphasizes the need for the development of new therapeutic strategies against GBS.

The current standard of care for GBS disease prevention and treatment is antibiotics. In the United States, pregnant individuals undergo routine antepartum screening via a rectovaginal swab^8^. GBS-positive mothers or mothers with undetermined GBS status are given intrapartum antibiotic prophylaxis to prevent vertical transmission^8^. However, there are serious and growing concerns about the sole reliance on antibiotics. First, antibiotics do not completely prevent GBS disease as intrapartum antibiotics do not impact late-onset GBS disease or recurrence of GBS disease in neonates^9^. Similarly, antibiotics also do not completely prevent recurrence of GBS after the end of GBS treatment for non-pregnant adults^10^. Second, while penicillin is the first-line antibiotic for intrapartum prophylaxis and for adult infection treatment^8^, second-line antibiotics used for those with beta lactam allergies are not as effective as penicillin in preventing disease or preventing GBS recurrence^11,12^, potentially posing a serious gap in effective care for those with beta lactam allergies. Concerningly, antibiotic resistance to penicillin and second line antibiotics is increasing^13,14^, putting all populations at risk.

Finally, there are mounting concerns about the overuse of antibiotics and their known adverse effects, especially in neonates. The use of antibiotics in early life is increasingly connected to short- and long-term adverse effects. Intrapartum antibiotics deplete the maternal vaginal microbiome. Mounting research underscores the importance of this early exposure for proper immune system development in part due to the microbiome^15^. Additionally, the use of intrapartum antibiotics is linked to marked changes in the infant microbiome up to a year after birth^16–18^. Notable changes include decreases in *Bifidobacterium* spp.^16,17,19–21^, a keystone species of the developing infant microbiome. As early life exposure to beneficial microbes is linked to the development of the immune system^22^, intrapartum antibiotics have potential adverse effects on the infant. In fact, children born to GBS-positive mothers have an increased risk of developing childhood asthma^23^.

This study develops a mouse model to begin to understand the understudied GI reservoir of GBS. Notably, GBS is asymptomatically carried in the GI tracts of 15-30% of adults^24^. The GI GBS population is a potential source for community spread^9,25,26^ and can predispose female reproductive tract GBS colonization^27^. Targeting GBS in the GI tract first requires a deeper understanding of GBS physiology in this environment. While parallels can be drawn between the GI tract and other body sites where GBS can cause disease, there has yet to be targeted investigation of asymptomatic GBS in the adult mammalian gut. Additionally, this is the first reported *in vivo* model to study GBS in the adult GI tract to our or knowledge. Using an *in vivo* transposon sequencing (Tn-Seq) screen, we have begun to identify GBS-encoded factors, such as capsular polysaccharide, that enable long-term GBS GI colonization based on. Additionally, this model establishes a versatile framework for future study into the bacterial and host factors that shape GBS persistence in the gut.

## Results

### Murine GI carriage of GBS strain COH1

We considered two important host-determined variables when establishing the model: diet and antibiotic use. Because diet is a key determinant of gut microbial ecology, we modeled a major feature of the Western lifestyle by feeding mice a fiber-deficient (FD) diet^28,29^ . Most adults in the United States fail to meet recommended daily fiber intake^30^, and this chronic deficiency alters both the microbiota and the metabolic environment of the gastrointestinal tract. Therefore, we rationalized that the FD diet allowed us to mimic the nutritional context in which GBS is likely to occur in the GI tract of adults in the United States and other parts of the industrialized world. In addition, antibiotic usage is not a risk factor for GBS carriage^24^, suggesting that GBS can invade human microbiomes in the absence of large-scale ecological disturbances. Therefore, we did not pre-treat mice with antibiotics prior to inoculation with GBS.

After inoculation of non-antibiotic treated mice fed the FD diet with GBS strain COH1, we observe long-term colonization of the murine GI tract for up to 2 months with fluctuations in overall GBS abundance, as determined by fecal sampling (**Figure 1**). Mice showed no behavioral signs of disease such as decreased activity, hunched posture, ruffled fur, or changes to feeding and drinking habits at any time point. Together, these observations are consistent with long-term asymptomatic carriage of GBS.

**Figure 1:**
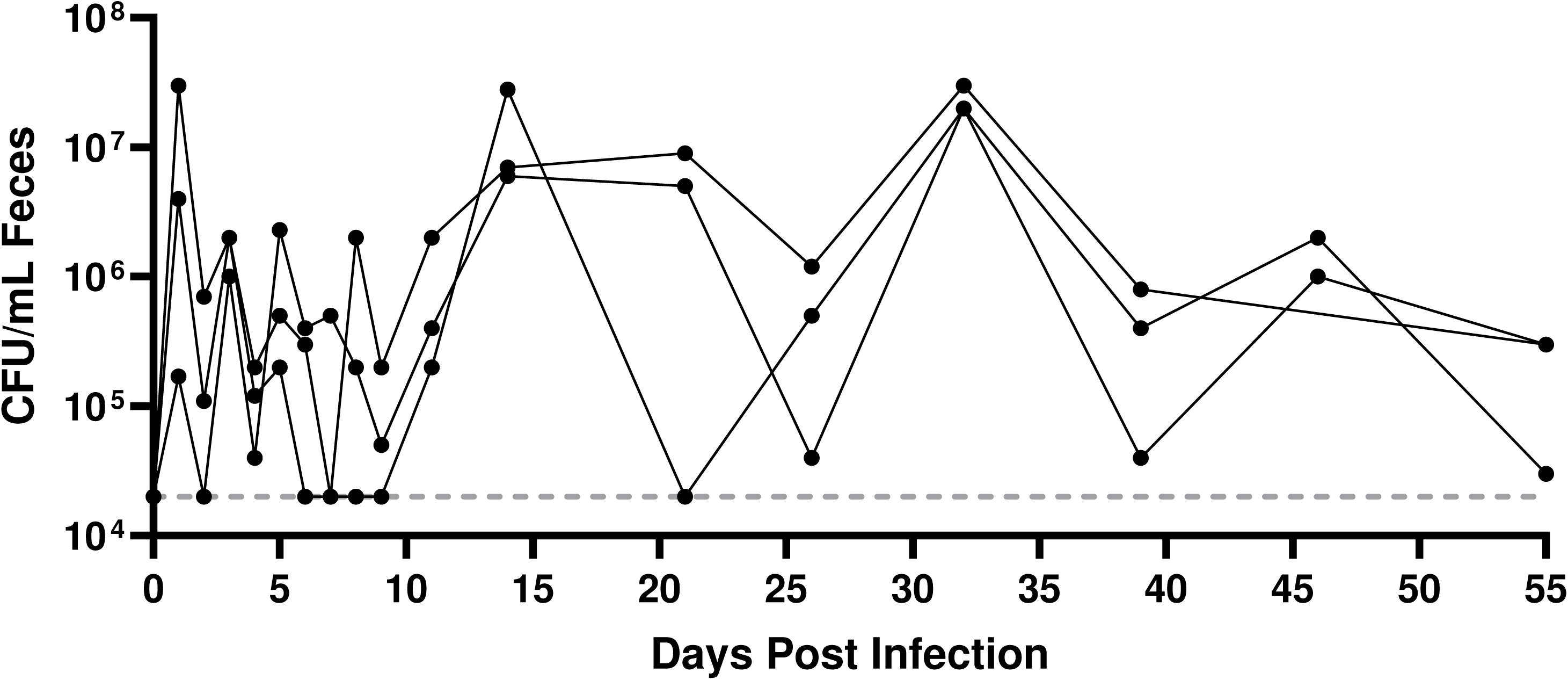
Long term gastrointestinal colonization of mice with GBS. Female C57BL/6 mice (n=3) were maintained on a fiber deficient diet for 1 week prior to inoculation with GBS strain COH1 via droplet feeding (see Methods). Feces were collected from the mice at the indicated time points and GBS was enumerated on Granada agar. Lines represent data collected from individual mice over time. The horizontal dotted line represents the limit of detection of the GBS plating assay (20,000 CFU/mL feces).

Because a common concern in murine GI models is that coprophagy could artificially maintain fecal burdens of microbes that poorly colonize, we housed mice individually on wire bottom cages to limit coprophagy and found no difference in GBS burdens compared to control mice that we co-housed under standard husbandry conditions (**Figure S1**), confirming that GBS GI colonization in the mice represents a stable population rather than repeated re-inoculation with transiently colonizing GBS. We also tested whether host biological sex influenced colonization and found no difference in GBS burdens between male and female mice (**Figure S2**), consistent with human survey data indicating similar GBS carriage between males and females^24^. These results establish that our model recapitulates robust, asymptomatic GBS carriage in the adult GI tract independent of coprophagy or host biological sex.

### Generalizability of the model to other GBS strains

GBS strains are commonly classified into ten recognized serotypes (Ia, Ib, II-IX)^31,32^ with serotype distribution in asymptomatic screening and invasive disease varying geographically^33–35^. In the US, the most common serotypes detected in human surveys are Ia, Ib, III, and V^7,36,37^. COH1 is a serotype III clinical isolate commonly used in laboratory research^38–40^ (**Figure 1**). To test the generalizability of GBS GI colonization to other strains commonly used in GBS research (which also mirror serotypes commonly detected in the US), we colonized mice with GBS strains COH1, A909 (serotype Ia)^39,41^, and two serotype V strains (CJB111)^42,43^ and the hypervirulent CNCTC 10/84^38,39^).

Like COH1, we observe that CJB111 and CNTC 10/84 can robustly colonize the GI tract over the length of the experiment while A909 has low GBS levels that decrease below the limit of detection within 4 days of inoculation (**Figure 2A**). Comparison of the average area under the curve (AUC) for fecal CFU counts further illustrates this difference and reinforces the differences in colonization between strains (**Figure 2B**). Together, these findings support the idea that genetic variation between GBS strains contributes to GBS fitness in the adult GI tract, as has been observed in the neonatal GI tract^44^.

**Figure 2:**
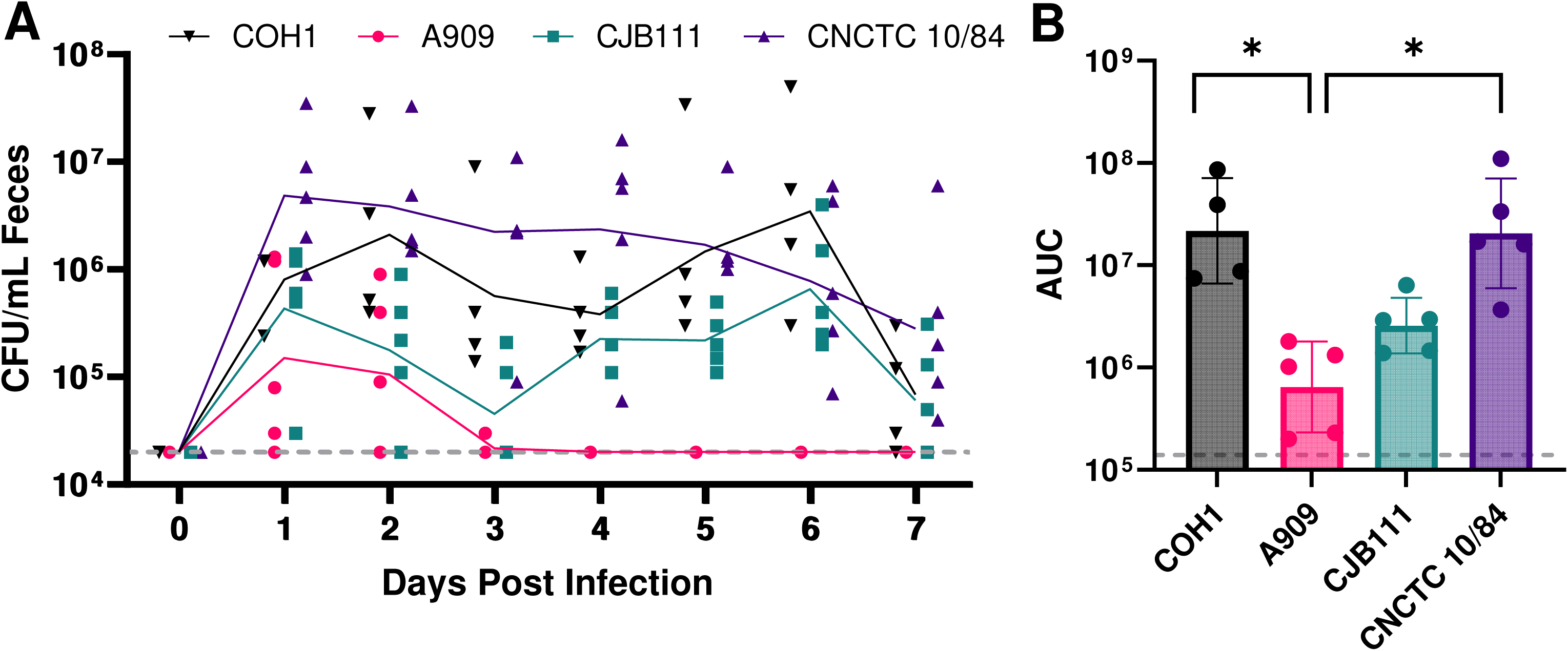
Gastrointestinal colonization with diverse GBS strains. Female C57BL/6 mice (n=4 for COH1, n=5 per group for all others) were maintained on a fiber deficient (FD) diet for 1 week prior to inoculation with GBS strains COH1, A909, CJB111, or CNCTC 10/84 via droplet feeding (see Methods). **(A)** Feces were collected from the mice at the indicated time points and GBS was enumerated on Granada agar. Points represent individual mice at each time point and line represents geometric mean of GBS burden per strain per day. The horizontal dotted line represents the limit of detection of the GBS plating assay (20,000 CFU/mL feces). **(B)** Area under the curve (AUC) from 0 to 7 days post infection. Points represent individual mice, bar represents geometric mean, and error bars represent geometric standard deviation. The horizontal dotted line represents the lower limit AUC (140,000). Statistical significance determined by Kruskal-Wallis test and Dunn’s multiple comparison test (*= p-value<0.05).

### Genome-wide analysis of GBS factors required for survival in the murine gastrointestinal tract

We next sought to identify which genetic factors GBS requires to colonize and survive in the murine GI tract. We utilized an existing GBS Tn mutant library^45–48^ in the CJB111 (Serotype V) background^42,43^ to colonize the murine GI tract. Three groups of mice (6 per group) were colonized with 1e7 CFU of the GBS Tn library in biological triplicate. Mice were monitored daily for signs of illness and fecal samples were collected daily (**Figure S3AB**). At 7 days post colonization, mice were euthanized and cecal content collected (**Figure S3C**). The input and cecum recovered libraries were processed as described in the materials and methods.

To identify transposon insertion sites, sequenced reads were mapped to the GBS CJB111 genome (GenBank Accession CP063198), which identified 1066 genes as significantly underrepresented (adj. p-value < 0.05, log2FC ≤ -2) and 12 genes as significantly overrepresented (adj. p-value < 0.05, log2FC ≥ 2) in the cecum compared to the input library (**Figure 3A** and **Table S1**). Due to the large number of significantly underrepresented genes from this initial analysis, we restricted further analysis to genes in the top 25^th^ percentile of significantly underrepresented genes (L2FC ≤ -6.46). These genes were assigned clusters of orthologous groups of proteins (COGs). COGs including carbohydrate transport and metabolism, amino acid transport and metabolism, nucleotide transport and metabolism, inorganic ion transport and metabolism, and cell wall/membrane/envelope biogenesis had the greatest number of genes in the top 25th percentile of significantly underrepresented genes (**Figure 3B**). We identified multiple genes known to contribute to GBS infection and adherence as significantly underrepresented such as genes that are involved in β-hemolysin/hemolytic pigment, two-component regulatory systems, pilus genes, as well as metal carbohydrate transport (**Table 1** and **Table S1**).

**Figure 3:**
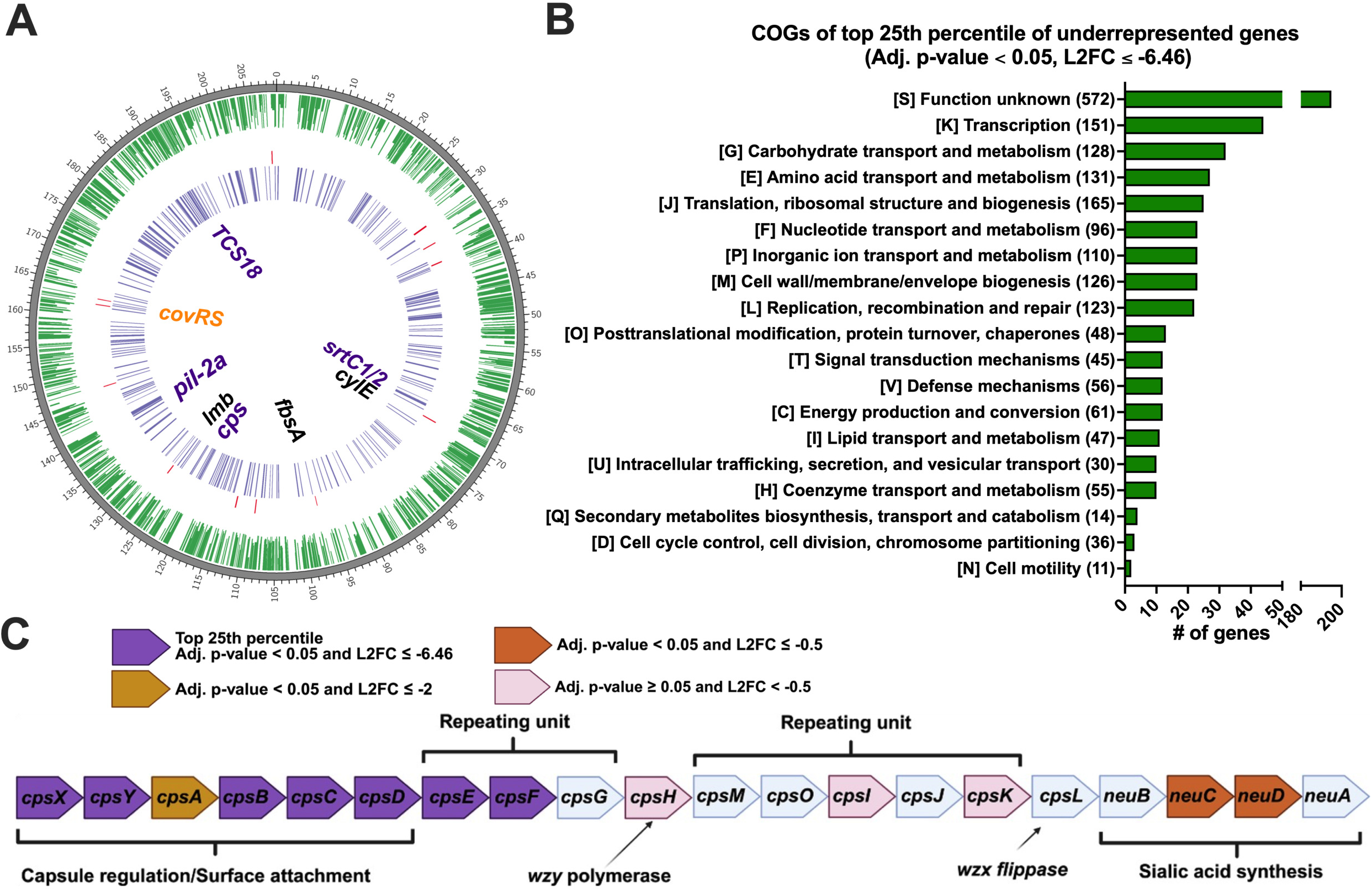
*In vivo* transposon-sequencing (Tn-Seq) of GBS CJB111 highlights genes possibly involved in GI colonization. **(A)** CIRCOS plot showing from outer ringer to inner ring; dark grey, genome length in Kb; green, statistically significant underrepresented genes (adj. p-value < 0.05 with Log2Fold Change (L2FC) ≤ -2) ; red, statistically significant overrepresented genes (adj. p-value < 0.05 with L2FC ≥ 2); purple, genes in the top 25^th^ percentile (adj. p-value < 0.05 with L2FC ≤ -6.46). Select genes are indicated in the center: top 25% of significantly underrepresented genes are labeled purple, other significantly underrepresented genes are labeled in black, and nonsignificant genes are labeled in orange. **(B)** COG distribution of the top 25% of significantly underrepresented genes. **(C)** . GBS Type V capsule biosynthesis gene operon.

**Table 1:** Underrepresented mutants of interest identified by *in vivo* gastrointestinal TN-seq.

| Open Reading Frame | Gene Name | Description | Log2FC |
| --- | --- | --- | --- |
| ID870_03480 | cpsY | LysR family transcriptional regulator | -8.15 |
| ID870_03510 | cpsF | UDP-N-acetylglucosamine--LPS N-acetylglucosamine transferase | -7.91 |
| ID870_03490 | cpsB | tyrosine-protein phosphatase | -7.73 |
| ID870_03495 | capA/cpsC | capsular polysaccharide biosynthesis protein | -7.38 |
| ID870_03500 | cpsD | tyrosine-protein kinase | -7.33 |
| ID870_00125 | phoU | phosphate signaling complex protein | -7.31 |
| ID870_00130 | TCS18_R | response regulator transcription factor | -7.84 |
| ID870_00135 | TCS18_S | two-component sensor histidine kinase | -7.33 |
| ID870_06015 | srtC2 | class C sortase | -7.46 |
| ID870_06020 | srtC1 | class C sortase | -7.26 |
| ID870_02610 | srtC4 | PI-2a pilus assembly sortase | -6.75 |
| ID870_02595 | pilA | PI-2a pilus adhesin | -6.74 |
| ID870_02600 | pilB | PI-2a pilus major subunit | -6.69 |
| ID870_02615 | pilC | PI-2a pilus subunit | -6.24 |
| ID870_05925 | cylE | cylE protein | -2.76 |
| ID870_03205 | lmb | metal ABC transporter substrate-binding lipoprotein/laminin-binding adhesin | -5.12 |
| ID870_04105 | fbsA | fibrinogen-binding adhesin | -6.14 |
| ID870_01600 | csrR/covR | response regulator transcription factor | -0.03 |

Among the most significantly underrepresented genes were genes of the capsular polysaccharide synthesis locus. The Type V capsule biosynthesis pathway of CJB111 is displayed in **Figure 3C**^49^. The genes, *cpsXYABCD*, are putative capsule regulation and surface attachment genes with 5 out of the 6 being in the top 25^th^ percentile of genes in the dataset (**Figure 3C**). Additional genes in the top 25^th^ percentile were *cpsEF,* putatively involved in the repeating unit biosynthesis (**Figure 3C**). Taken as a whole, this suggests the importance of capsule in GI survival, where the loss of any step in the biosynthesis pathway leads to decreased GBS survival.

### Loss of capsule leads to decreased GBS survival in the adult mouse GI tract

The Tn-seq experiment identified mutants in the capsule biosynthesis locus as being significantly underrepresented in the strain library present in cecal contents. To test the importance of capsule in GBS GI colonization, the colonization experiment was repeated using a CJB111 Δ*cpsD* mutant which has reduced abundance of capsule compared to WT ^45^. Daily fecal sampling shows the capsule mutant is less fit in the GI tract, with the majority of Δ*cpsD*-colonized mice having GBS burdens below the limit of detection by the end of the sampling period (**Figure 4A**). Comparison of the average area under the curve (AUC) for fecal CFU counts from mice colonized with CJB111 or CJB111 Δ*cpsD* further reinforces the role that capsule plays in colonization of the murine adult GI tract (**Figure 4B**). Taken together, these observations support that capsule is important in maintaining colonization of the adult GI tract.

**Figure 4.**
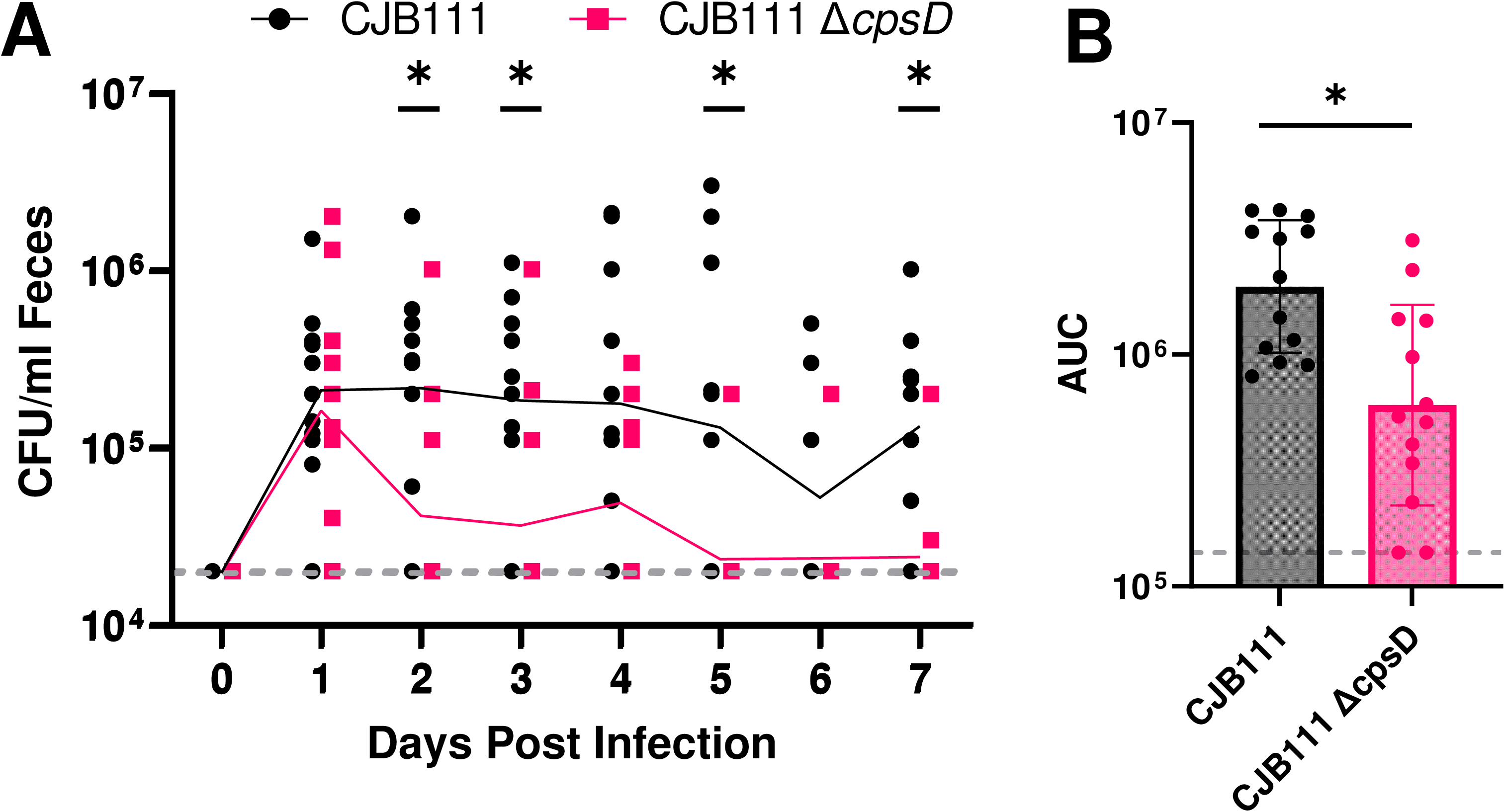
Capsular polysaccharide is important for GBS colonization of the adult mouse GI tract. Male and female C57BL/6 mice (n=13 per group) were maintained on a fiber deficient (FD) diet for 1 week prior to inoculation with GBS strains CJB111 WT or CJB111 Δ*cpsD* via droplet feeding (see Methods). **(A)** Feces were collected from the mice at the indicated time points and GBS was enumerated on Granada agar. Points represent individual mice at each time point and line represents geometric mean of GBS burden. The horizontal dotted line represents the limit of detection of the GBS plating assay (20,000 CFU/mL feces). **(B)** Area under the curve (AUC) from 0 to 7 days post infection. Points represent individual mice, bars represent geometric mean, and error bars represent geometric standard deviation. The horizontal dotted line represents the lower limit AUC (140,000). Mann-Whitney test (*= p-value<0.05) showed statistically discernible differences between AUC of the strains.

## Discussion

Group B *Streptococcus* is a serious health concern for humans across their lifespan. While current antibiotic-based interventions and treatments have been effective, the rise in antibiotic resistance and increasing awareness of the negative impacts of antibiotic-mediated microbiome disruption emphasizes the need for non-antibiotic approaches for mitigating GBS disease. In addition, the majority of invasive GBS diseases have been in non-pregnant adults in recent years. Unlike neonates and pregnant people, there does not exist a standard prophylaxis approach for non-pregnant individuals who predominantly acquire GBS by community spread. More research is required to understand the community reservoir of GBS, the adult GI tract.

Here, we present a robust model of GBS adult GI colonization that paves the way for a better understanding of this pathogen in this important but understudied niche and established that capsular polysaccharide is an important GBS-encoded molecular factor required for colonization. This contributes to a growing body of literature that highlights the importance of capsule for GBS to colonize and cause disease in a variety of host-associated niches^44,50,51^. We observed differences in the fitness of different GBS strains, with the Ia serotype strain A909 not colonizing well compared to serotype III and V strains that we tested (**Figure 2**). This is in contrast to previous neonatal experiments that showed the serotype Ia capsule positively affects neonatal GI colonization and infection relative to the serotype III capsule^44^. This could suggest differences in the role GBS capsule plays in the immature neonatal gut versus the adult gut. Further supporting the importance of capsule in GI colonization, we showed via an established TN-seq assay and our mouse model that multiple GBS capsule biosynthesis genes were in the top 25% percentile of underrepresented genes (Figure 3). We then showed that a serotype V capsule mutant is unable to colonize the GI tract *in vivo* to the same extent as the WT (Figure 4).

Other genes significantly underrepresented in the output library include the sortases *srtC1* and *srtC2*, known adherence factors (such as PI-2a^52^ and *lmb*^53,54^) that promote colonization and adherence to epithelial cells and extracellular matrix component. Notably, the CovR/S transcriptional regulation system, which has previously been identified as important in colonization and virulence^55,56^, was not significantly underrepresented. Instead, the two-component system TCS18 was in the top 25% of underrepresented COGs. TCS18, i.e. PhoBR, seems to impact GBS biofilm formation and hemolysin production by binding to the promoter regions of *hylA* and *ciaR*^57^. Taken together, this suggests that same genetic factors that enable colonization and virulence in other body sites could also be involved in establishing asymptomatic colonization in the GI tract.

A limitation of this study is that we focused solely on the large intestine (cecal contents and feces) and do not investigate GBS in other parts of the GI tract. This was done due to the large intestine having the largest amount of biomass in the GI tract. However, it is well known that the ecology of the microbiome varies widely down the length of the GI tract. *Streptococcal* species are hallmarks of the oral cavity and the small intestine^58,59^. While GBS has been isolated in the oral cavity^60^, it is not the majority member. This could suggest that GBS is outcompeted by other *Streptococcal* species in these niches, making GBS preferred GI niche the large intestine. Future work will leverage this model to establish GBS biogeography throughout the longitude of the GI tract and to identify the microbial and host factors important for maintaining colonization in these diverse locations.

This work also lays the groundwork for further study into how diet impacts GBS in the GI. The GI tract is a mosaic of fluctuating nutrients with host diet dictating availability and by extension, impacting bacterial fitness. GBS, like other *Streptococcus* species, has a reduced genome of approximately 2 Mb. As a result, GBS is an auxotroph for many essential compounds and depends predominantly on scavenging from the environment and other bacterial community members. We anticipate that this model will enable focused research into how host diet could be leveraged as a non-invasive, non-antibiotic-based intervention to decrease GBS GI colonization rates and mitigate subsequent disease.

We anticipate that this model will also enable future study into how GBS transmits from the GI reservoir to other body sites, such as the urogenital tract, and to other at-risk individuals, such as neonates and immunocompromised adults. GBS adult GI colonization predisposes human urogenital tract colonization^27^, and this model can be leveraged to explore the bacterial and host factors that drive this transmission. Additionally, longitudinal rectovaginal screening of pregnant people suggests GBS status can be transient^61^, suggesting transient factors such as host behavior and diet could impact GBS burdens and risk of transmission. This is echoed in our own findings where we see day-to-day fluctuations in GBS burdens. Future work could aim to address the host, and bacterial factors drive this variability.

The adult GI tract is an understudied but vital reservoir for GBS. A better understanding of GBS physiology in the GI tract is crucial in the development of targeted interventions. We have established a robust adult mouse model that recapitulates key human host factors such as being sex-independent and not requiring antibiotic pre-treatment. This model opens the door to a mechanistic dissection of how GBS persists in the gut and how this impacts carriage at other body sites, transmission to neonates, and subsequent neonatal and adult disease.

## Materials and Methods

### Bacterial strains and culture conditions

GBS strains COH1 (serotype III), A909 (serotype Ia)^39,41^, CJB111 (serotype V)^42,43^, CNCTC 10/84 (serotype V)^38,39^, and CJB111 Δ*cpsD*^45^ were maintained at -80°C as 25% glycerol stocks. Strains were routinely cultured on tryptic soy broth agar (TSB, Neogen) and incubated aerobically at 37°C. After overnight growth, a single colony was picked into 5 mL of TSB and was grown aerobically at 37°C for 16-24 hours. Liquid cultures were used as inocula for mouse experiments, described below.

### Murine model of GBS GI colonization

All animal studies were carried out in strict accordance with the University of Wisconsin-Madison Institutional Animal Care and Use Committee (IACUC) guidelines (Protocol #M006305). Six-to-eight week old C57BL/6 mice were purchased from Taconic and used as in-house breeders. In-house bred mice between 6 and 15 weeks of age were used in experiments. Mice were fed a fiber-deficient (FD) diet (Inotiv TD.150689) diet starting one week before GBS inoculation.

Mice were inoculated via oral droplet feeding method. This method was developed based on an existing murine model of *Klebsiella pneumoniae* gastrointestinal colonization ^62^. In brief, on the day of inoculation, food and water were withheld from mice 4 hours before oral droplet feeding. To prepare the inoculum, 5 mL overnight GBS liquid culture was centrifuged at 4,000 x g for 15 min. The resulting bacterial pellet was resuspended and washed once with phosphate-buffered saline (PBS), and resuspended in equal volume of sterile 2% sucrose. Using this inoculum, mice were fed ∼1e8 CFU/mL total of GBS, split between two 50-uL doses an hour apart via micropipette tip.

Feces were collected from mice directly into a microcentrifuge tube and were kept on ice until plating, upon which they were stored at -80°C. To quantify GBS burdens, 1 µL of fecal sample was resuspended in 200 µL PBS in sterile polystyrene 96-well tissue culture plates. Tenfold serial dilutions were prepared and 10 µL of each dilution was spread on Granada agar^63^ (a differential and selective medium for GBS) with two technical replicates. Granada agar plates were incubated anaerobically for 24 hours. Colonies were quantified and the technical replicates were averaged to determine GBS burden (limit of detection =2e4 CFU/mL).

### Murine GI colonization Tn-seq

Triplicate 1mL frozen aliquots of the pooled CJB111 pKrmit transposon library^47^ were thawed and resuspended in 4mL TSB with kanamycin at 300 µg/mL and grown overnight at 37°C in. Cultures were washed in PBS and then normalized to 1e8 CFU/mL in 2% sucrose in sterile PBS as described above. Mice were then fed the library via oral droplet feeding as described above. 100 uL of the input libraries were spread on CHROMagar Strep B with 300 µg/mL kanamycin in triplicate and incubated overnight at 37°C. Burdens of GBS were monitored via fecal sampling as described above with the exception that fecal samples were plated on CHROMagar Strep B with 300 µg/mL kanamycin. 7 days post infection, mice were euthanized and cecal content was collected. Cecal content was resuspended in 1 mL sterile PBS and spread on CHROMagar Strep B with 300 µg/mL kanamycin in triplicate and incubated overnight at 37°C to collect recovered transposon mutants. Bacterial growth from spread plates were collected using a sterile tissue scraper and the 6 mice per library pooled together. The pooled recovered transposon mutants were centrifuged at 4,000 x g for 15 min. Supernatant was then removed and bacterial pellets were then frozen and stored at - 80C before genomic DNA was extracted using a phenol-chloroform method^64^.

### Transposon library sequencing

Libraries were prepared and sequenced at the University of Minnesota Genomics Center (UMGC) according to https://www.protocols.io/view/transposon-insertion-sequencing-tn-seq-library-pre-rm7vzn6d5vx1/v1. Briefly, genomic DNA was enzymatically fragmented, and adapters added using the NEB Ultra II FS kit (New England Biolabs), and ∼50 ng of fragmented adapted gDNA was used as a template for enrichment by PCR (16 cycles) for the transposon insertions using mariner-specific (TCGTCGGCAGCGTCAGATGTGTATAAGAGACAGCCGGGGACTTATCATCCAACC) and Illumina P7 primers. The enriched PCR products were diluted to 1ng/ul and 10 ul was used as a template for an indexing PCR (9 cycles) using Nextera_R1 (iP5) and Nextera_R2 (iP7) primers. Sequencing was performed using 150 base paired-end format on an Element Aviti system to generate ∼40 million reads per library.

### Tn-seq bioinformatics analyses

Bioinformatic analyses were performed as previously described with minor modifications^45,46^. TRANSIT2 (v 1.1.7)^65^ was used to trim reads and align them to the CJB111 genome (CP063198) for analysis of transposon insertion sites. The Transit PreProcessor processed reads using default parameters with the Sassetti protocol, primer sequence ACTTATCAGCCAACCTGTTA, and mapped them to the genome using Burrows-Wheeler Alignment (BWA)^66^. Insertion sites were normalized using the beta-geometric correction (BGC) in TRANSIT2 and analyzed using the site-restricted resampling was performed using default parameters, with the addition of ignoring TA sites within 5% of the 5’ and 3’ end of the gene, to compare the insertion counts recovered from the cecum vs the input library. All sequencing reads have been deposited into NCBI SRA under BioProject accession: PRJNA1328725

### Statistical analysis

All statistical analysis, except those done for the Tn-seq (see above), were performed using GraphPad Prism 10. Specific statistical analyses are noted in the relevant figure legends.

## Acknowledgments

This material is based upon work supported by the National Institutes of Health (R35GM150996 to AJH, R01AI153332 to KSD), the National Science Foundation Graduate Research Fellowship Program (DGE-2137424 to JL), and the Robert H. and Carol L. Deibel Distinguished Graduate Fellowship in Probiotic Research (JL) and the American Heart Association grant 23POST1013835 (LRJ), Any opinions, findings, and conclusions or recommendations expressed in this material are those of the author(s) and do not necessarily reflect the views of the funders. AJH is the Judy L. and Sal A. Troia Professor in Gastrointestinal Disease Research at the University of Wisconsin-Madison.

JL and LRJ performed experiments, analyzed the data, and prepared the display items. KSD and AJH provided key insights, tools, and reagents. JL, LRJ, and AJH wrote the paper. All authors approved the manuscript prior to submission.

## Conflict of interest

The authors declare no conflict of interest.

